# IS-capades of *Klebsiella pneumoniae*: Insertion sequences drive metabolic loss in obscure sub-lineages

**DOI:** 10.1101/2025.07.24.666535

**Authors:** Ben Vezina, Claire White, Helena B. Cooper, Kathryn E. Holt, Jane Hawkey, Kelly L. Wyres, Margaret M. C. Lam

## Abstract

**Introduction:** *Klebsiella pneumoniae* is an opportunistic pathogen which causes a wide spectrum of infections within healthcare settings and the community. Four *K. pneumoniae* sub-lineages defined with cgMLST/LINcodes are known to cause distinct infections of the nasal and/or upper respiratory passages: SL91 and SL10031 (also referred to as subspecies *ozaenae*), SL10032 (subspecies *rhinoscleromatis*) and SL82. These sub-lineages have also demonstrated reduced carbon source utilisation, which in other species has been linked with high loads of insertion sequences (IS).

**Methods:** We performed comparative genomics, analysed IS and constructed genome-scale metabolic models for available public sequences from these four sub-lineages and compared them to other sub-lineages from the wider *K. pneumoniae* population.

**Results:** The four focal sub-lineages displayed significantly higher IS loads (median range 88 to 120 per genome) compared to other *K. pneumoniae* sub-lineages (median range 12 to 73). Notably, each *K. pneumoniae* sub-lineage had unique IS profiles, consistent with distinct evolutionary trajectories of IS acquisition and expansion. Across sub-lineages, higher IS loads were inversely associated with the number of metabolic model genes per genome (R^2^ = 0.16, p <0.001), as well as predicted aerobic substrate utilisation for phosphorus sources (R^2^ = 0.39, p<0.001) as per a second-degree polynomial regression model (n = 1,664 genomes). Additionally, the four IS-dense sub-lineages displayed a combination of convergent sub-lineage-specific substrate utilisation losses including parallel loss of 3-Phospho-D-glycerate, D-Glycerate-2-phosphate, Phosphoenolpyruvate utilisation as carbon/phosphorus sources. Finally, inspection of IS insertion sites demonstrated frequent and non-destructive insertion next to transcriptional, carbohydrate and amino acid metabolism genes.

**Conclusions:** IS loads were significantly and inversely associated with metabolic substrate usage within *K. pneumoniae*, whereby sub-lineages that had higher numbers of IS also had reduced metabolic capacity. We hypothesise that an insertional tolerance model explains these findings, whereby IS can only insert into “metabolically-tolerable” sites for the individual cell and any impacts on metabolism are not detrimental for survival.

## Introduction

*Klebsiella pneumoniae* is an opportunistic pathogen which causes a diverse range of infections within healthcare settings such as pneumonia, bloodstream, surgical site and urinary tract infections. It is also a causative agent of infections in the community, associated with a unique subset of *K. pneumoniae* sub-lineages distinct from those that cause healthcare-associated infections (1). As a species, *K. pneumoniae* exhibits a remarkable amount of genome diversity, with hundreds of unique ‘deep-branching’ sub-lineages and significant variation in its accessory genome content (1, 2). This diversity is likely an important driver of the different lifestyles and variable virulence profiles of *K. pneumoniae*.

Two sub-lineages of *K. pneumoniae*, clonal groups CG67 and CG90/91 based on 7-gene MLST and formally designated as subspecies *rhinoscleromatis* and *ozaenae*, respectively, cause rare but unique chronic infections typically associated with the nasal and/or upper respiratory passages. Whole genome sequencing has since clarified that both are divergent sub-lineages of *K. pneumoniae* rather than separate subspecies as previously characterised **Error! Reference source not found**.(3). The *K. pneumoniae* LINcode taxonomic scheme (based on a 629-loci core genome MLST) assigns *K. pneumoniae* into >705 discrete sub-lineages (4). Our sub-lineages of interest are SL10032 (previously *rhinoscleromatis* or CG67) and SL10031/SL91 (previously *ozeanae* or CG90/CG91). SL10032 strains cause a chronic granulomatous disease described as rhinoscleroma, whereas SL91 and SL10031 typically cause atrophic rhinitis or ozena. There have been at least 63 cases of *rhinoscleromatis* (5-9) and 45 *ozaenae* (8-22) infections documented in modern medical literature dating back to 1944, although there appears to be archaeological evidence of rhinoscleroma as far back as 300-600 AD from Maya (modern day Guatemala) (23). While these sub-lineages are usually associated with the nasopharyngeal sites, there is strong historical proof that they are able to colonise other body sites including the urinary tract, soft tissue, blood, cerebral spinal fluid and gastrointestinal tract (11, 12, 18, 21, 24, 25). Additionally, these sub-lineages have also been found in non-human hosts, isolated from cockroaches, cattle and chicken meat (13, 21, 22).

Aside from their characteristic clinical presentations, these sub-lineages display a reduced metabolic capacity in biochemical tests compared to other *K. pneumoniae* sub-lineages (3, 26), explaining why they were originally considered distinct sub-species. These sub-lineages, along with SL82 (ST82 or CG82 based on 7-gene MLST), have demonstrated reduced phenotypic carbon source utilisation (3). A link between this metabolic reduction and niche/pathogenic lifestyle adaptation has been posited (3). This phenomenon has also been observed in other bacterial pathogens such as *Shigella* species and *Bordetella pertussis* (27). Shigella in particular is associated with the accumulation of insertion sequences (IS) (28). Reduced carbon utilisation is also seen in SL82 (3), which along with *ozaenae* were unique amongst *K. pneumoniae* in lacking the *mrkD* type III fimbrial adhesin. SL82 expressed the virulence-associated K1 capsule serotype. Similar to the other sub- lineages of interest, SL82 was noted to be strongly associated with respiratory infections (8/11 BioSample IDs with sample metadata).

In this study, we performed a systematic analysis on a dereplicated collection of 1,664 completed and 210 draft *K. pneumoniae* genomes, to investigate the prevalence and impacts of IS on metabolic capacity in these four sub-lineages of interest. We hypothesised that, similar to *Shigella*, IS caused genome degradation and the loss of metabolic capabilities in these sub-lineages.

## Methods

### Genome acquisition, assembly and annotation

Complete *K. pneumoniae* genome assemblies (n=2,302) were initially downloaded from NCBI RefSeq (accessed on 04/07/2024, **Table S1**). To further expand genome numbers and improve robustness of population-level inferences, 210 dereplicated NCBI BioSample IDs of target sub-lineages were obtained from BIGSdb (29) by selectively downloading genomes which matched the 4 sub-lineages of interest and their associated Sequence Types when no LINcode was determined, including SL10032 (ST67, ST3818, ST3819), SL91 (ST91, ST381, ST3766, ST3768), SL10031 (ST91) and SL82 (ST82, ST3764). Where available, short-read sequence reads were downloaded from SRA (n=201), otherwise assembled sequences were downloaded from Genbank (n=20) (accessed on 29/10/2024, **Table S1**).

Short-read sequence data were assembled with Unicycler v0.5.1 (30). Low quality genomes were removed if they had >300 graphical fragment assembly dead ends, or in absence of assembly graphs, an N50 of <65,000. This threshold was more lenient than previously defined quality control metrics (31), to account for IS fragmenting short read draft assemblies, resulting in larger contig numbers and dead end counts. Remaining genomes (n=2,108) were then dereplicated using Assembly-Dereplicator v0.1.0 (32) using the following specifications: ‘--threshold 0.0003’. This resulted in a collective dataset of 1,874 dereplicated genomes. All genomes were annotated using Bakta v1.8.1 (33).

### Lineage assignment and genotyping

Dereplicated genomes were uploaded to Pathogenwatch (https://pathogen.watch/) for genotyping; namely to confirm species and assign sub-lineage, clonal groups and LIN codes (4, 34). A neighbour-joining tree was generated with PopPUNK v2.4.0 (35) as follows. The ‘create-db’ function was used with the following options: ‘--sketch-size 1000000 --min-k 15 -- max-k 29 --qc-filter prune’. The ‘fit-model’ function was subsequently used with the following options: ‘bgmm --ranks 1,2,3,5 --graph-weights --K 3’. Additionally, the ‘poppunk_visualise’ function was used, with the ‘--distances’ and ‘--previous-clustering’ parameters, to output a neighbour joining tree. Kaptive v3.1.0 (36, 37) was used to identify capsule synthesis loci (KL), and genotyping was performed with Kleborate v3.1.3 (38).

### Metabolic model construction

Metabolic models were constructed using Bactabolize v1.0.3 (31) with the *Kp*SC pan v2.0.1 model (39) and the --draft_model command. Growth across 1,278 conditions was simulated using the --fba command on M9 minimal media, where positive growth was defined at a biomass threshold of ≥0.0001 as previously described (40).

### Insertion Sequence analysis

Plasmids were filtered out from the complete genome sequences using seqtk v1.3 (41), with the following command: “seq -L 4500000”. Genomes were rotated to *dnaA* at base pair position 1 using rotate v1.0 (42) with the following command: “-s gtgtcactttcgctttggcagcagtgtcttgcccgattgcaggatgagtt -m 5”, to facilitate comparison of chromosomal synteny. Rotated, plasmid-free genomes (i.e. the chromosome) were used for the remaining analyses.

ISEScan v1.7.2.3 (43) was used to identify IS elements, which also identifies novel IS elements with divergence from known IS in the ISEScan database. For comparisons within SL82, SKA v1.0.0 (44) was used to determine single nucleotide variants (SNV) using ska fasta followed by ska distance -s 25. Within sub-lineages, specific phylogenetic minimum evolution trees were constructed using SNV distances inferred from SKA, optimized via Nearest Neighbour Interchange (NNI) with the fastme.bal command from R package ape v5.8 (45), then midpoint rooted via midpoint.root from phytools v2.3-0 (46). All-vs-all whole genome alignments were performed using MUMmer4 v4.0.0 (47) using “nucmer -p”, then coordinates extracted using “show-coords -d -l -T”.

Genome locus comparisons were performed by extracting gene coordinates via slice_multi_gbk.py (https://github.com/bananabenana/slice_multi_gbk) and Clinker v0.0.31 (48) was used for visualisation.

### Statistical analysis and code

R v4.4 (49) and RStudio v2024.04.2+764 (50) were used for statistical analysis and visualisation. R packages tidyverse v2.0,0 (51), ggtree v3.12.0 (52), colorspace v2.1-0 (53), ggpmisc v0.6.0 (54), ggpubr v0.6.0 (55), ggh4x v0.2.8 (56), rstatix v0.7.2 (57), gggenomes v1.0.0 (58) and patchwork v1.2.0 (59) were used. All R code can be found at Figshare (doi: 10.6084/m9.figshare.28341917).

Linear regression was performed between IS loads and the ratio of nucleotides assigned to IS vs total chromosomal sequence. IS loads between sub-lineages were compared using a Kruskal-Wallis test with Holm adjustment for multiple comparisons, followed by a Dunn’s *post hoc* test with Holm adjustment for multiple comparisons.

## Results

### Dataset description

To explore the impact of IS in *K. pneumoniae*, we utilised two dereplicated datasets: i) a collection of complete genomes (n=1,664) used to quantify the exact numbers of IS within chromosomal sequences and phylogenetic clusters; and ii) an expanded collective dataset (n=1,874) which included targeted inclusion of draft assemblies. Throughout the manuscript, we will explicitly state which dataset was used. We found that 26 draft assemblies failed Pathogenwatch’s quality control measures (**Table S1**), as they had >500 contigs, despite having high levels of genome completeness (≤300 graphical fragment assembly dead ends). We suspected high contig numbers may be caused by large numbers of IS and these assemblies were kept for analysis.

Significant diversity was observed in the expanded dataset (n=1,874), which was made up of 244 sub-lineages (based on cgMLST). To ensure appropriate population-level inferences, only sub-lineages with n≥3 genomes were included for further analysis, resulting in a dataset of n=1,441 genomes representing 65 sub-lineages. The majority of genomes were from sub-lineages associated with known multi-drug resistant (n=1,013 across 14 MDR SLs; 61.69%) or hypervirulent clones (n=129 from 4 hypervirulent SLs; 7.86%) as defined by Wyres *et al* (1) (**Table S1**).

### Some sub-lineages display unusually high IS loads

Plasmid-free complete genomes (n=1,664) were screened for IS, and the numbers and types (i.e. IS families) were compared across the 65 sub-lineages with n≥3 genomes (**Fig. 1**). As expected, linear regression indicated a strong, positive correlation (R^2^ = 0.89, p<0.001) between IS loads and the ratio of nucleotides assigned to IS vs total chromosomal sequence (**Fig. S1**). Across each sub-lineage, the number of IS per genome (i.e. IS-load) varied from 10 to 131 (median 31). A total of 132 unique IS from 23 distinct IS families were detected across this dataset. Sub-lineages contained 3 to 20 distinct IS families, of which only three IS families (IS*3*, IS*21* and IS*NCY*) were found in every sub-lineage. Lineage-specific IS patterns were identified (**Fig. S2, Table S2**). Aside from five novel IS (Novel 20, 24, 189, 272 and 348, which were detected in sub-lineages SL43, SL91, SL107, SL147 and SL17, respectively. No other IS families were restricted to a single sub-lineage. An additional 19 IS were rarely detected across the population (i.e. found in <10 sub-lineages).

**Fig. 1:**
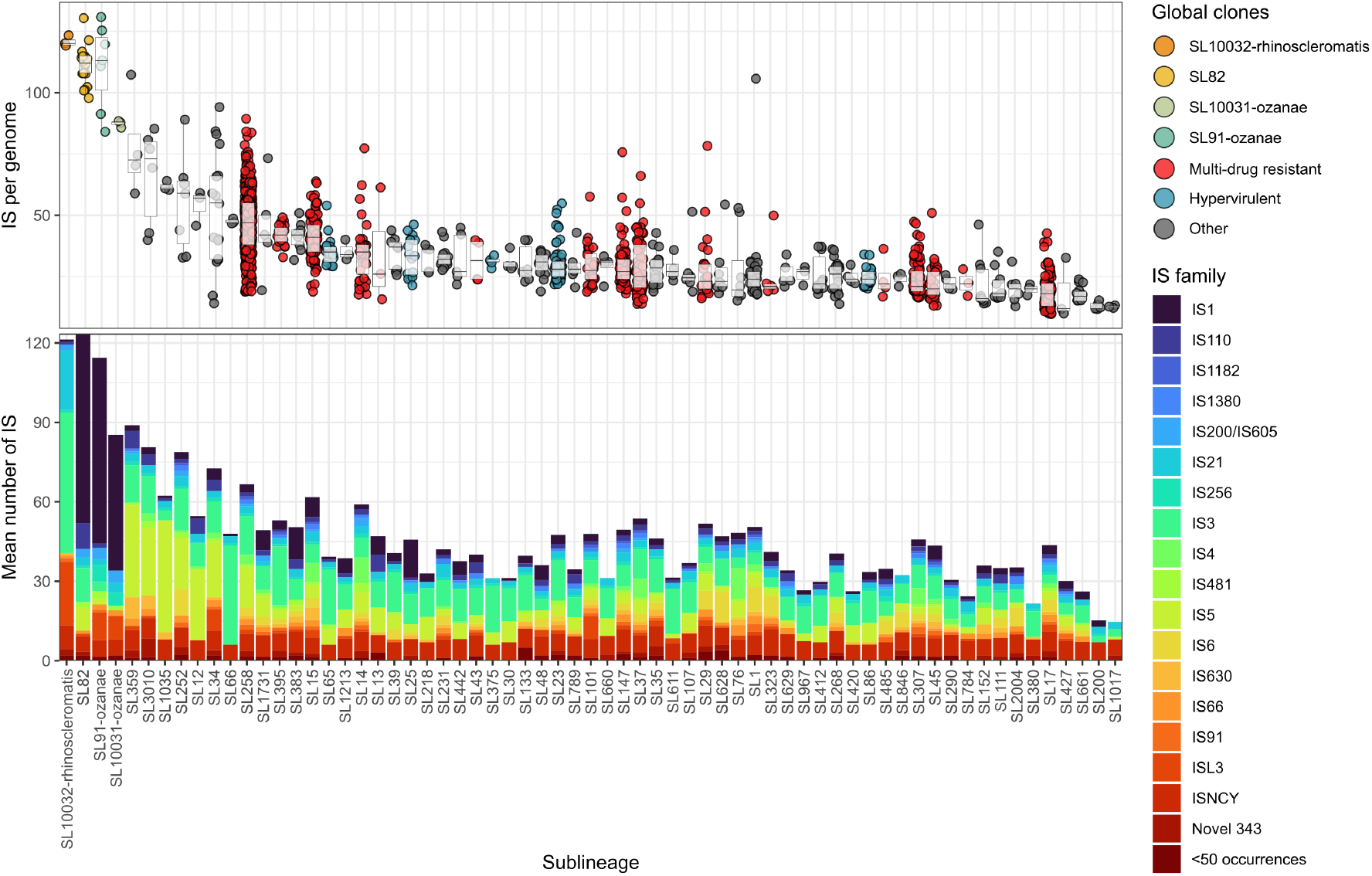
Sub-lineages display significantly different IS loads. Distribution of Insertion Sequences (IS) within 65 *K. pneumoniae* sub-lineages containing n≥3 complete genomes (n=1,441). The top panel shows the IS per genome while bottom panel shows the breakdown of IS families. Sub-lineages are coloured by their relevant global clone information (60), where each point represents a single genome. ‘<50 occurrences’ refers to IS families which were not shown in this figure due to their low abundance. Significance not shown for brevity - raw data and results of Dunn’s post *hoc test* in **Table S2**.

Sub-lineages SL82, SL91-ozaenae, SL10031-ozaenae and SL10032-rhinoscleromatis carried the highest IS loads, with median 88 to 120 IS per genome compared to other SLs (median 12 to 73, **Fig. 1**). Kruskal-Wallis with Holm error correction indicated sub-lineages had significantly different IS loads (p < 0.0001). This was followed by a Dunn’s *post hoc* test with Holm error correction, showing SL82 had significantly more IS per genome than 37 other sub-lineages (p-values <0.05), while SL91-ozaenae, SL10031-ozaenae and SL10032-rhinoscleromatis had significantly higher IS loads than 25, 5 and 6 other sub-lineages, respectively (**Table S2**). There were no significant differences between SL0032-rhinoscleromatis (median 120 ± 2 IQR), SL91-ozaenae (median 113 ± 21.5 IQR), SL82 (median 112 ± 7 IQR) or SL10031-ozanae (median 88 ± 1). We focused on these four IS-dense sub-lineages for the remaining analyses. The complete genome set was supplemented with additional short-read assemblies that could be assigned LINcodes, resulting in a total of n=1,874 genomes, which included n=4 SL10032-rhinoscleromatis, n=15 SL91-ozaenae, n=5 SL10031-ozaenae, and n=36 SL82.

This statistical analysis also revealed that several other sub-lineages displayed significantly higher IS loads compared to others (**Table S2**). This included genomes that corresponded to MDR (SL258, SL15, SL395), hypervirulent (SL65) and common clones (SL34, SL3010), although there appeared to be within-clone variation in many sub-lineages. For example, the number of IS per genome within sub-lineage SL258 ranged from 22 to 80. Within sub-lineage variability in IS loads was typically larger for sub-lineages represented by more genomes. With the exception of our four sub-lineages of interest, the IS loads were generally similar across other clones regardless of MDR or hypervirulence status.

Aside from isolates of unknown sampling site (n=42, 70% of genomes from the four sub-lineages of interest), the remaining isolates were recovered from human nasopharyngeal sites including sputum (n=8), nasopharynx (n=6), maxillary sinuses (n=2), pleural cavity (n=1) and throat (n=1) (**Table S1**). One additional isolate was isolated from blood (n=1). While source metadata was missing for many of the SL10032-rhinoscleromatis and SL91-ozanae isolates, sampling dates indicated collection between 1920 to 1952. Many samples collected during this period were generally of clinical origin. While there is likely a sequencing bias for clinical isolates, these sub-lineages have previously been isolated from a variety of non-human sources including chicken meat, cattle and cockroaches (13, 21, 22), although no such isolates are currently represented in public genome repositories.

### IS profiles varied within IS-dense sub-lineages

The four focal IS-dense sub-lineages contained 16 (SL91-ozaenae), 15 (SL82), 13 (SL10031-ozaenae) and 10 (SL10032-rhinoscleromatis) IS families (**Fig. 2**). Of these, seven IS families (IS*1*, IS*L3*, IS*21*, IS*66*, IS*256*, IS*NCY* and IS*200*/IS*605*) were detected across all four lineages (**Fig. S2**). The IS profile of SL10032-rhinoscleromatis was the most distinct from the others while the closely-related *ozaenae* sub-lineages and SL82 shared highly similar IS profiles, consistent with their divergence from a more recent common ancestor.

**Fig. 2:**
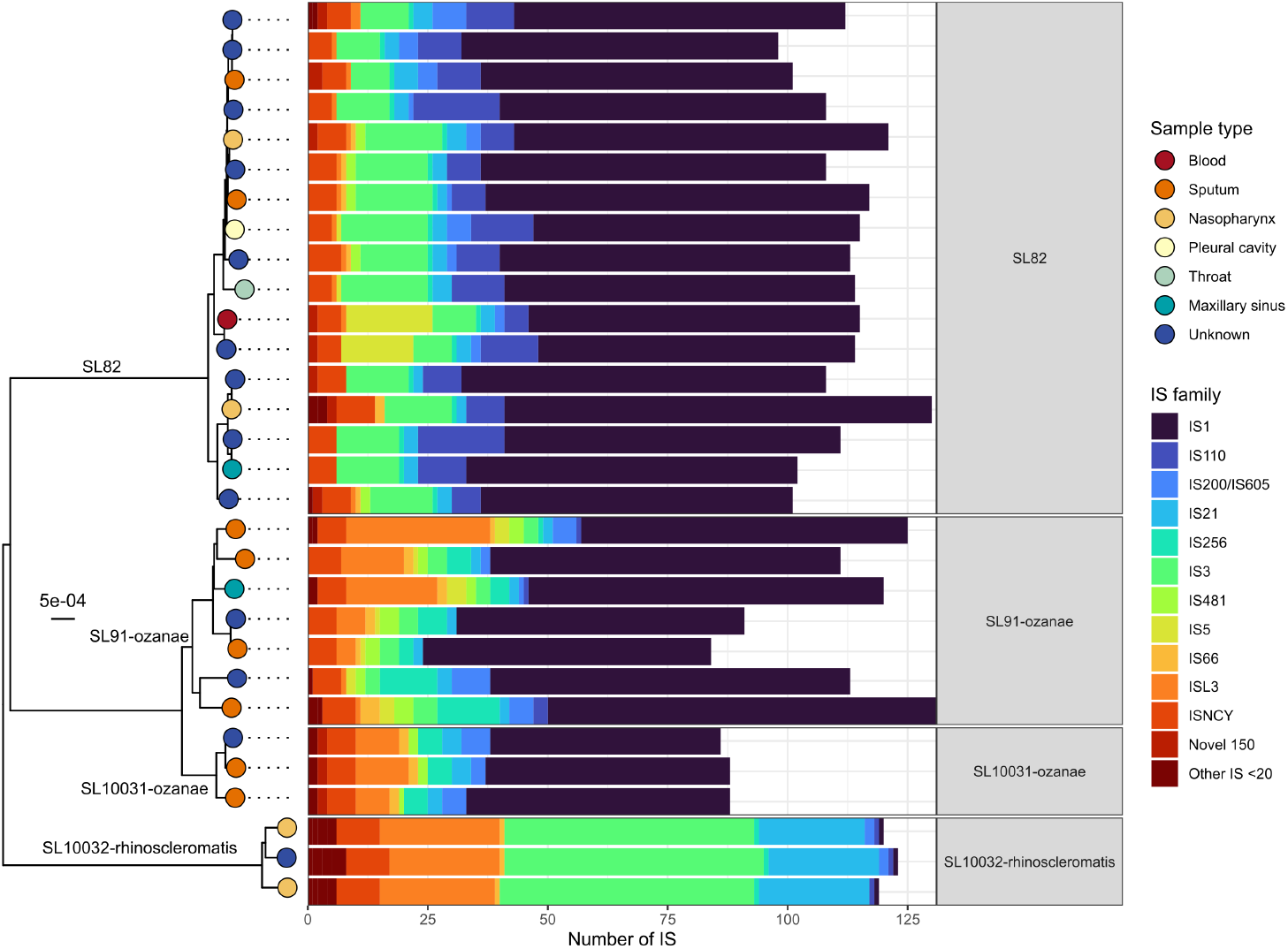
IS family prevalence differ between *K. pneumoniae* sub-lineages. Distribution of Insertion Sequence (IS) families across the four most IS-dense sub-lineages, aligned against a neighbour-joining tree generated from core k-mers, whereby each tip represents a unique genome and is coloured by the sample type. Only complete genomes with plasmids filtered out were used in this analysis.

Most notable is the considerable expansation of IS*1* in SL82 (mean 71.53±6.31 SD copies per genome), SL91-ozaenae (70.14±7.9) and SL10031-ozaenae (51.3±3.5) genomes, while it was conversely rarer in SL10032-rhinoscleromatis; (1±0). IS*1* expansion (mean ≥10 copies) was not specific to these sub-lineage and was found in six others including hypervirulent SL25 and other clones SL10022 and SL383.

There were, however, notable differences between the four focal sub-lineages and which IS families had undergone significant expansions. For example, expansion of IS*3* was observed in SL10032-rhinoscleromatis (mean of 53±1 SD copies per genome) and SL82 (12.8±3.2), while SL91-ozanae and SL10031-ozanae carried 3.7±1 or no IS*3*, respectively. IS*L3* was highly expanded in SL10032-rhinoscleromatis (24±1), SL91-ozaenae (10.57±10.81), SL10031-ozaenae (9±2), but was largely absent in SL82 (1±0). In SL10032-rhinoscleromatis, IS*3*, IS*L3* and IS*21* comprised the largest proportions of IS (mean 53±1, 24±1 and 22.7±0.6 SD copies per genome, respectively), while carrying fewer IS*1* (mean 1±0 copies).

### IS load impacts strain metabolism and redundancy

We next analysed the impact of IS on metabolism across the entire collective dataset (complete and short read-assembled genomes), although IS loads of short-read assemblies were not used in correlative measures. Metabolic models were built for all sub-lineages with n≥3 isolates, resulting in 1,874 genomes across 70 sub-lineages. Of these, 9 sub-lineages required minimal gap filling to simulate growth on M9 media + D-glucose (mean 2.89±2.33 SD gap-filled reactions), with SL34 (n=4) being the most prevalent lineage, caused by poorer draft genome assemblies. This is a standard process which improves simulated growth prediction accuracy (31).

Our model predictions were able to simulate 61/99 previously generated biochemical tests (3) and models are unable simulate the remaining substrates. As such, we compared metabolic modelling growth predictions to these 61 phenotypic substrates to generate accuracy statistics. As many of the biochemically-tested isolates do not have corresponding genome data, we compared model accuracy on a per-sub-lineage rather than individual basis to account for this (40). F1 scores varied, ranging from 0.72 to 0.83 (**Table 1**). The inaccuracies were largely driven by false positives, where the models predicted growth due to presence of intact genes, while phenotypic tests predicted no growth, as seen by the high recall values ranging from 0.88 to 0.94.

**Table 1:** Summary of accuracy metrics of simulated metabolic model growth predictions. A total of 61 compatible phenotypic biochemical tests from (3) were used. Full data and confusion matrix can be found in **Table S3**.

| Sub-lineage | Accuracy | Precision | Recall | Specificity | F1 score | Biochemical test N | Metabolic model N |
| --- | --- | --- | --- | --- | --- | --- | --- |
| SL82 | 0.62 | 0.60 | 0.88 | 0.32 | 0.72 | 5 | 5 |
| SL91-ozaeanae | 0.66 | 0.65 | 0.89 | 0.32 | 0.75 | 11 | 4 |
| SL10031-ozaeanae | 0.66 | 0.63 | 0.94 | 0.30 | 0.75 | 9 | 36 |
| SL10032-rhinoscleromatis | 0.75 | 0.76 | 0.93 | 0.43 | 0.83 | 4 | 15 |

We next examined the impact of IS load on strain metabolism. Analysis of only the completed genomes demonstrated a clear inverse relationship between IS load per genome and metabolic capacity, whereby high-IS sub-lineages displayed reduced metabolism (**Fig. 3**). Fitting a squared polynomial to the data, the number of model genes was inversely associated with IS load (R^2^=0.16, p <0.001) and slightly less with model reactions (R^2^=0.16, p <0.001). For substrate usage, only aerobic use of phosphorus sources displayed a moderate inverse relationship (R^2^=0.39, p <0.001). The association between IS loads and reduced anaerobic usage of substrates was weaker compared to aerobic substrate usage, likely due to the reduced metabolic capacity of *K. pneumoniae* as a whole under anaerobic conditions (40).

**Fig. 3:**
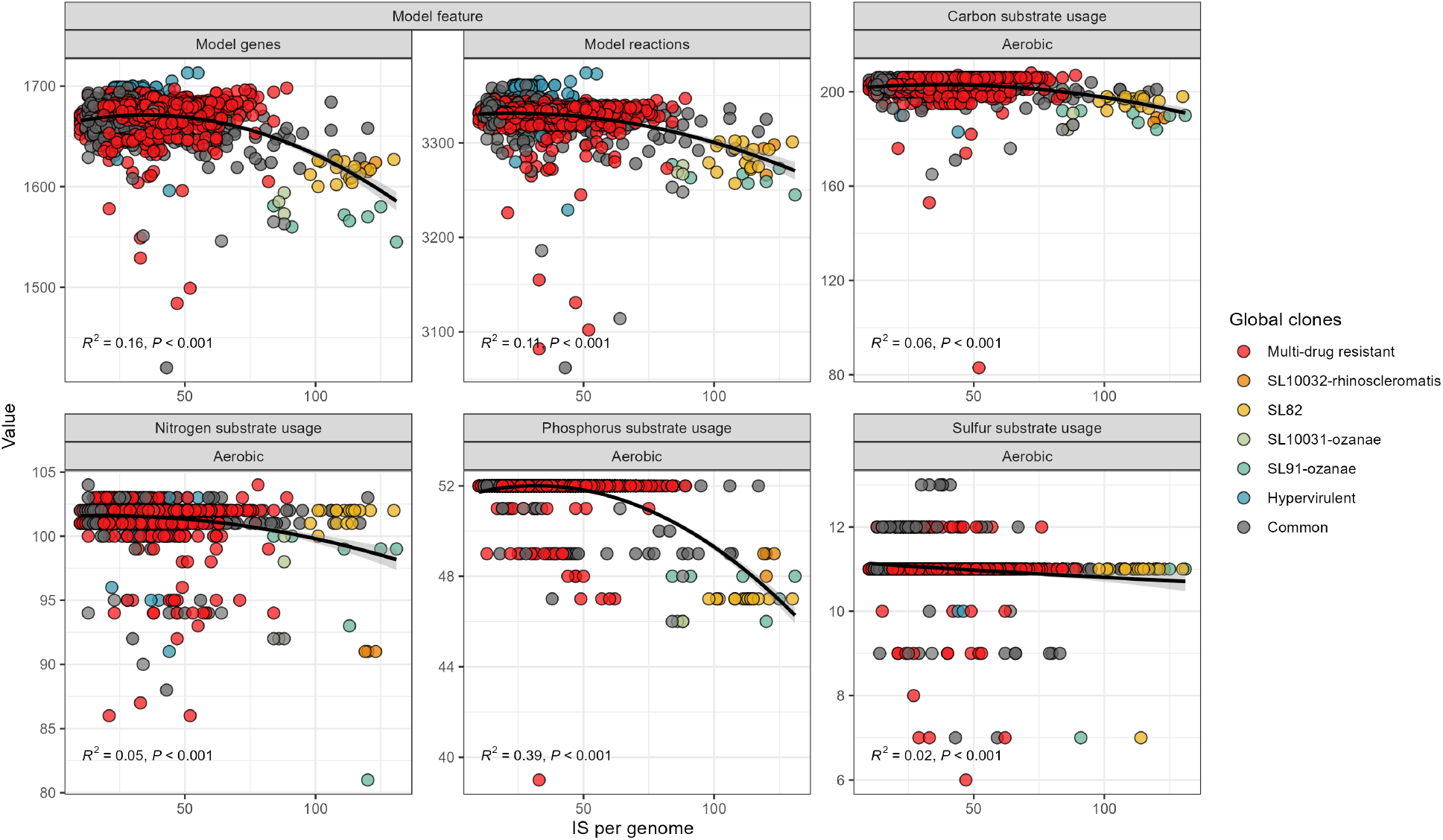
Number of IS per genome is inversely related to loss of metabolic substrate usage. Number of insertion sequences (IS) per genome plotted against key metabolic model metrics, with a squared polynomial model fitted to data. Each point represents a completed genome.

The metabolism of some over-represented sub-lineages such as MDR global clone SL258 (n=427) were unaffected by IS load (R^2^<0.01, p <0.263 to <0.96) despite a wide IS range (19–89) (**Fig. S3**). To control for this over-representation effect, we calculated mean values with each sub-lineage and re-examined the relationships (**Fig. S4**). Increasing IS load was now weakly associated with reduced aerobic use of carbon (R^2^=0.23, p <0.001), nitrogen (R^2^=0.13, p <0.001) and moderately associated with phosphorus (R^2^=0.42, p <0.001) sources using a squared polynomial model. The reduced metabolic capacity of sub-lineages with higher IS-loads was consistent with the similar inverse relationships of total number of metabolic model genes (R^2^=0.31, p<0.001) and model reactions (R^2^=0.25, p<0.001). There was no correlation between IS load and the number of pseudogenes (R^2^ < 0.01, P = 0.775), which was also previously reported in *Shigella* species (28).

A total of 232 reference model genes were missing across the complete genomes from SL82, SL91-ozaenae, SL10031-ozaenae and SL10032-rhinoscleromatis. These were a combination of core and variable genes across the dataset generally. Flanking sequences of up to 50bp around IS sites were matched against missing model genes to determine if IS had interrupted them directly. Only five genes across 16 total occurrences showed evidence of direct IS interruption. This meant that 1.5% of missing model genes can be directly attributed to a known IS insertion. For example, the loss of *kpnE*, an essential component of the KpnEF spermidine antiporter was absent in a single SL82 genome (accession GCF_900452625, coordinates 2591281:2597710 on contig 1), caused by insertion of IS*1* (**Fig. 4**).

**Fig. 4:**
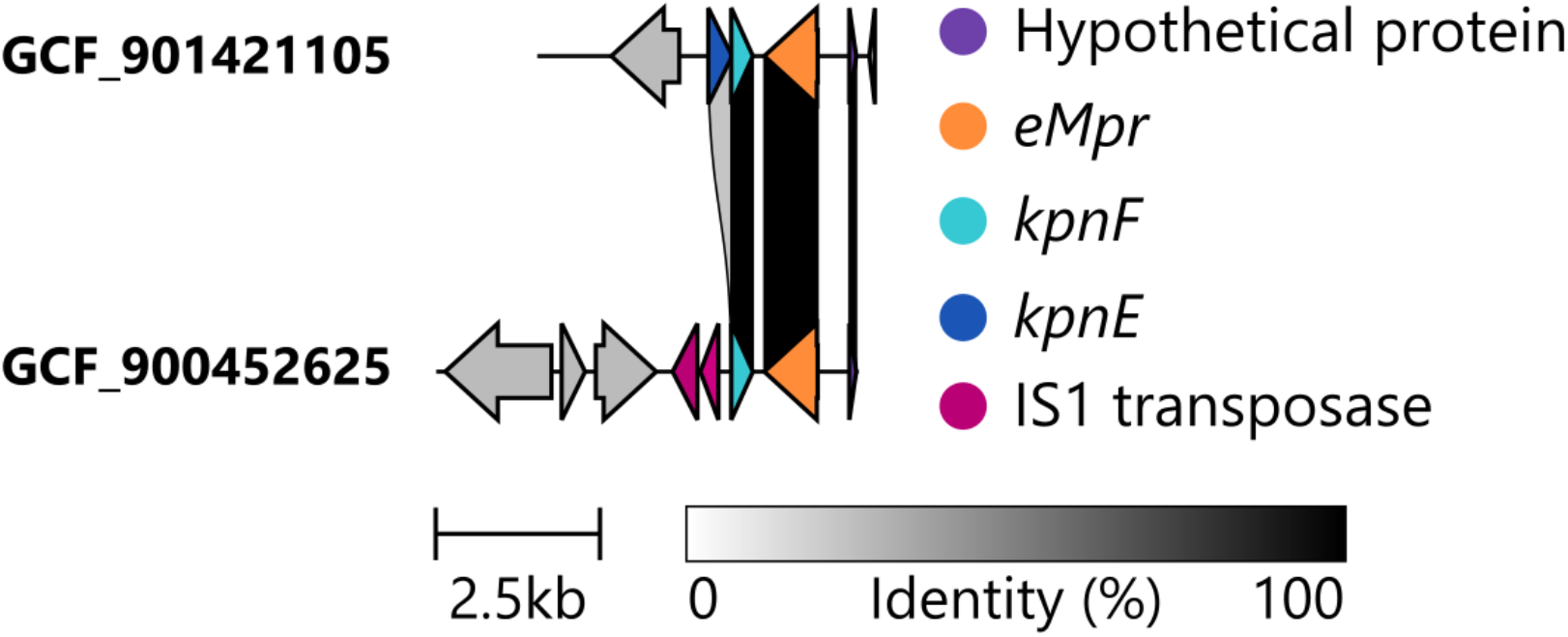
IS*1* interruption of the *kpnE* gene required for the KpnEF spermidine antiporter in an SL82 genome (accession GCF_900452625.1).

### Metabolic evolutionary parallelism

Aside from their loss of total substrate usage, these sub-lineages displayed distinct but convergent metabolic trajectories. Previously, 616 core metabolic traits were found to be conserved across the 48 most prevalent *K. pneumoniae* sub-lineages from a dataset comprising large-scale studies (4,621 genomes; n≥15 genomes in each) (40). By raw counts, each of the genomes from the four focal sub-lineages in our analysis appear to have completely lost the ability to utilise 15 to 20 core *K. pneumoniae* substrate growth conditions, while intermediately losing 12 to 49 (**Fig. 5, Table S3**).

**Fig. 5:**
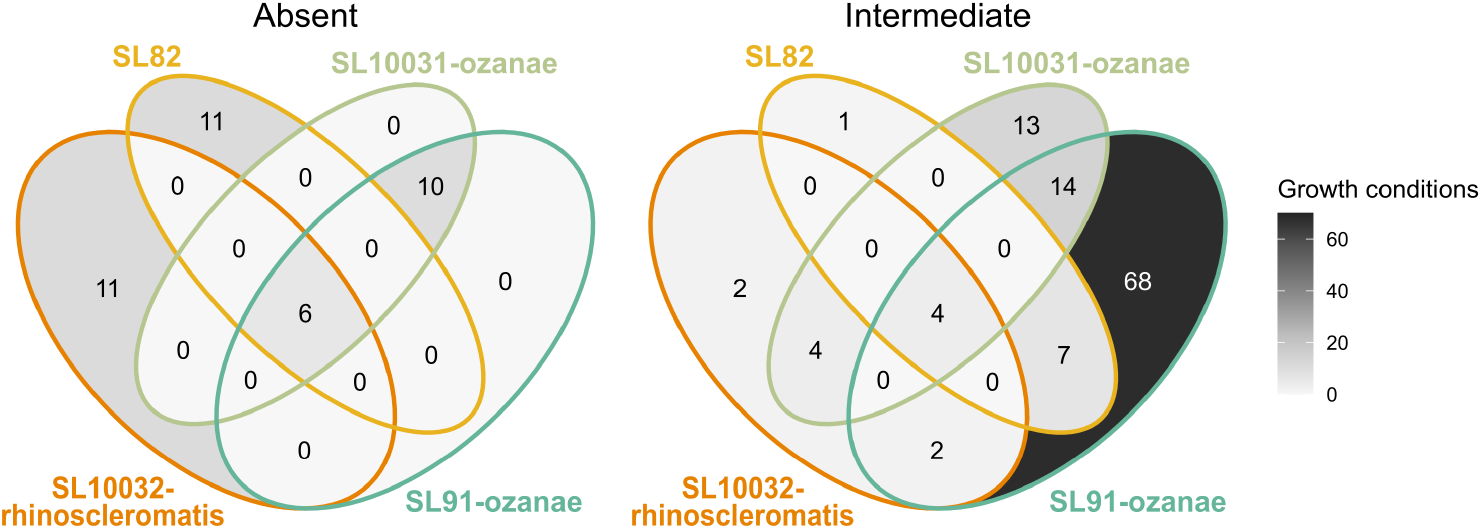
Sub-lineages display unique and convergent metabolic losses. Venn diagrams showing number of *K. pneumoniae* core substrate usages intermediately (usage within >0% & <95%) or completely lost (usage within 0%) within sub-lineages.

There was evidence of convergent substrate utilisation losses whereby all four sub-lineages have lost the ability to use 3-Phospho-D-glycerate, D-Glycerate-2-phosphate, Phosphoenolpyruvate as carbon or phosphorus sources. This was due to the loss of a periplasm antiporter *pgtP* and associated operon *pgtABC*, responsible for the import of these substrates from the periplasm to the cell. Due to the complete contextual loss of this operon and numerous intra-genomic rearrangements, it is not clear if this loss was IS-mediated. As expected based on their genetic relatedness, SL91-ozaenae and SL10031-ozaenae displayed 10 conserved substrate losses, likely as a result of common ancestry. Despite similar IS profiles between SL82 and the two ozaenae sub-lineages, they displayed little overlap of metabolic losses. Sub-lineage-specific complete metabolic losses included SL82’s loss of 3-hydroxyphenylacetic acid usage as a carbon source, SL10032-rhinoscleromatis’s loss of 5-Aminopentanoate, L-Lysine and 4-Aminobutanoate usage as carbon and nitrogen sources and SL91-ozaenae/SL10031-ozaenae’s loss of (R)-3-Hydroxybutanoate, Cytosine and Myo-Inositol hexakisphosphate usage as carbon and nitrogen sources (**Table S3**).

Siderophore virulence genes associated with iron uptake were common within these sub-lineages, with 29/60 genomes displaying Kleborate virulence scores ≥3 (i.e. carry yersiniabactin and aerobactin) (38) (**Table S1**). Yersiniabactin was found in 3/4 SL10032-rhinoscleromatis genomes (lineage: *ybt* 11), 13/15 SL91-ozaenae (lineage: *ybt* 10), 2/5 SL10031-ozanae (lineage: *ybt* 16), although was truncated in 12/18 cases across this dataset. Aerobactin was found in all SL10032-rhinoscleromatis (lineage: *iuc* 4), 6/15 SL91-ozaenae (lineage: *iuc* 2A), 2/5 SL10031-ozaenae (lineage: *iuc* 2A) and 17/36 SL82 (lineage: *iuc* 2A), with only one truncated case in SL10032-rhinoscleromatis. Salmochelin (encoded by *iro*) was found only in 17/36 SL82. Previously reported *mrkD* (type 3 fimbriae adhesin) (3) was completely absent in all focal sub-lineages with the exception of SL10032-rhinoscleromatis (present in 5/5 genomes). The *rmpADC* locus which controls capsule expression and hypermucoviscosity was also commonly detected in genomes from these four sub-lineages. It was found in 32/36 SL82 genomes (lineage: *rmp* 2A, truncated in 26 instances), all SL10032-rhinoscleromatis genomes (lineage: *rmp* 4, truncated in 1 instance), and 1/15 SL91-ozanae (lineage: *rmp* 2A, truncated). These virulence gene truncations were associated with the ends of assembly contigs, likely caused by IS insertions.

All SL82, SL10031-ozaenae and SL10032-rhinoscleromatis isolates were typeable and displayed homogenous capsular loci (KL1, KL5 and KL3, respectively), consistent with previous reports (3) based on capsule typing. In contrast, SL91-ozaenae isolates displayed varying K loci with closest matches to KL4 (n=13), KL6 (n=1), KL1 (n=1), with all but the KL6 genome being untypeable – again due to capsule gene truncations and fragmented assemblies, likely caused by IS insertions within the capsule loci, which has been previously reported (63). As public assemblies lacked reads, it was not possible to leverage assembly graphs to determine which IS caused contig breaks – though in complete genomes such as GCF_014218685.1, IS*1* was responsible for the capsule null predictions.

### IS variation within sub-lineages confers strain-level metabolic diversity

To study the localised impact of IS further, we analysed all available SL82 genomes (n=17) as this sub-lineage had the largest number of complete genomes. After re-orientating the genomes to the same starting position, we mapped IS across them and inspected IS insertion sites and chromosomal synteny (**Fig. 6**). While many of these genomes within this sub-lineage shared similar IS profiles and IS density, no two genomes were identical with respect to IS insertions, highlighting the dynamic nature of IS and consequent losses or gains of DNA segments. Additionally, chromosomal inversions and rearrangements were common, which were often flanked by IS, reasonably explaining the rearrangements occurring at homologous, repeat sites.

**Fig. 6:**
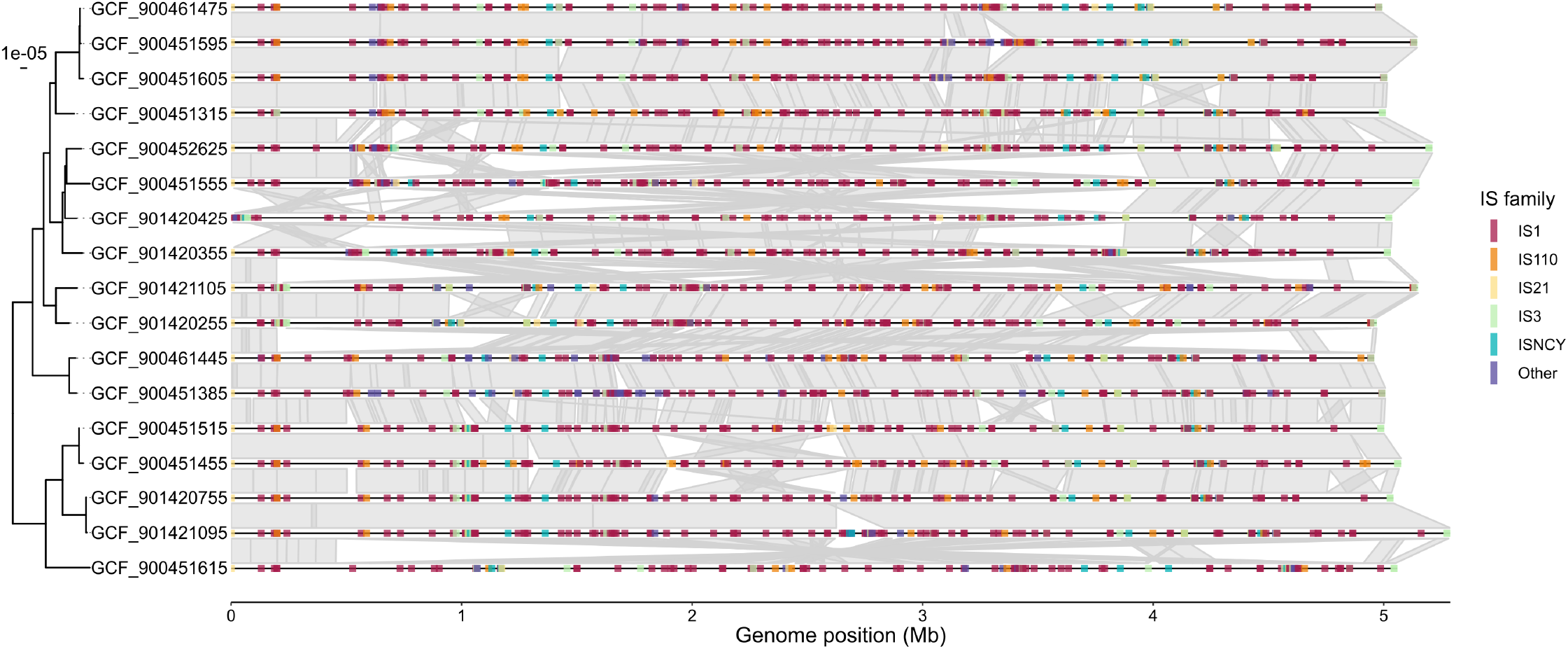
Insertion Sequences vary locationally within SL82. Comparison of insertion sequences (IS) and homologous blocks between SL82 complete chromosomes. Tree shown is a SNV distance tree showing 17 genomes. Grey links between genomes represent homologous sequence blocks as aligned by nucmer with standard parameters. Coloured lines show IS family. All genomes start at *dnaA*.

The genomes within SL82 were also predicted to have small but varying predicted aerobic metabolism across 24 substrates, with 13/17 genomes showing loss of usage for at least one substrate. The most variable substrates were 4-Hydroxybenzoate (usage observed in 7/17 genomes), then L-alanine-D-glutamate-meso-2,6-diaminoheptanedioate (13/17 genomes), and L-Galactonate (15/17 genomes). The remaining 21 substrates were absent only in a single genome (16/17).

## Discussion

Reduced metabolic versatility has previously been documented in select sub-lineages of *K. pneumoniae*: SL10032-rhinoscleromatis, SL10031-ozaenae, SL91-ozaenae and SL82 (3). This is thought to be an important driver of their unique ecological and pathogenic profiles, particularly as these strains are often isolated from the upper respiratory tract but are by no means restricted to these sites or human host (13, 21, 22). In this study, application of genome-scale metabolic modelling supported significant loss of substrate usage across SL82, SL10031-ozaenae, SL91-ozaenae and SL10032-rhinoscleromatis compared to other *K. pneumoniae* sub-lineages. In particular, these sub-lineages appear to have lost the ability to utilise gut microbe/human metabolites for energy including 3-Phospho-D-glycerate, D-glycerate-2-phosphate, phosphoenolpyruvate, 3-hydroxyphenylacetic acid and 5-aminopentanoate (64-68), along with myo-Inositol hexakisphosphate found in plant tissues and human diet (70) and human metabolites (R)-3-hydroxybutanoate (liver) (71) and 4-Aminobutanoate (neurotransmitter) (72). These hint that these sub-lineages have deviated from other *K. pneumoniae* sub-lineages that readily colonise the gut prior to infection (69).

We demonstrate for the first time that these four sub-lineages also have significantly higher IS loads (**Fig. 1**), which has been linked to streamlining of metabolic profiles in *Bordetella pertussis* and various *Shigella* species (27, 28). IS has been previously quantified within *K. pneumoniae* SL258 (ST258) (73), which found a stable repertoire of IS elements and small within-lineage variation (specifically, an increase in IS load in one subclade of SL258, referred to as ST258B). This is consistent with the IS profiles and within-sub-lineage variation that were observed in our study (**Fig. 1**, **Fig. 2**, **Table S2**), such as intermediate rather than complete loss of core *K. pneumoniae* substrates within a sub-lineage (**Fig. 5**). Notably, the lack of impact IS had on the metabolism of SL258 differed from that of the four focal sub-lineages. This indicated that *K. pneumoniae* sub-lineages have different relationships and IS-mediated evolutionary trajectories. This analysis was limited by the sampling biases of public data, and the relative rarity of these particular sub-lineages.

Direct IS interruptions of metabolic genes were detected in only a few instances (only 5/323 examples were found). We hypothesize that IS may have been responsible for the initial disruption of other key metabolic genes, whose loss was selectively tolerated at the time of transposition, leading to subsequent genetic degradation of the associated metabolic loci over time. The inverse relationship between IS loads and substrate usage, primarily aerobic carbon and phosphorus usage (**Fig. 3, Fig. S4**) is consistent with this hypothesis. Insertion of an IS may have caused the initial interruption of the gene but have been subsequently purged from the genome over time -either via recombination, rearrangement or genome degradation. Additionally, higher IS loads potentiate a higher rate of genomic rearrangement (74), deleting genes without direct insertion. This highlighted that DNA sequence analysis of isolated pathogens cannot capture the *in vivo* conditions or genetic history which shaped the organism even when historical isolates are used. Hawkey *et al*. noted significant genome degradation from IS insertions in *Shigella* species which caused metabolic usage losses (28). We did not observe any correlation between IS load and genome length in our dataset, contrasting to *Shigella* where the two appeared to be related. The metabolic gene losses in both *Shigella* and the four *K. pneumoniae* sub-lineages of interest provide insight into the cellular processes required for these pathogens to colonise and progress to invasive human disease. Our data show that while these sub-lineages displayed loss of some *K. pneumoniae*-specific core traits, they retained use of 552 to 587 core metabolic traits. Given conservation of these core metabolic traits across all *K. pneumoniae* sub-lineages, these are likely key to the species’ ability to colonise and/or cause infections.

As it currently stands, the metabolic models have broad agreement with previously published biochemical tests (3, 26) (**Table 1**) whereby these IS-dense sub-lineages exhibited considerable loss of metabolic potential (**Fig. 5**). The decaying, polynomial relationship between increased number of IS per genome and loss of substrate growth conditions, as well as metabolic genes and reactions (**Fig. 3**), demonstrates this effect. Lower accuracy scores were observed, (**Table 1**) and can be explained by various reasons. False positives are not always indicative of model inaccuracies, but usually indicate the presence of metabolic genes which failed to be expressed in the experimental condition due to gene regulation (40, 75), or phenotypically-tested isolates do not have intact copies of key genes. Discrepancies likely stem from strain choice used in biochemical tests, which may not be reflective of the larger population. In some cases, false-positives could be explained by IS interrupting the promoter/regulatory regions required for this expression to occur. For example, the genes required for L-histidine, L-tyrosine, ethanolamine, quinate, 2-Ketoglutarate and putrescine utilisation are all present in the genomes but phenotypically showed no growth. Alternatively, if these false positives did not arise from gene regulation issues, there may be incorrect assumptions being made in the metabolic models, such as over-assignment of metabolic genes. A metabolic model gene is considered ‘present’ if there is ≥25% bi-directional coverage (standard value) (31, 39, 76). This would indicate our model-based analysis is likely underrepresenting the impact of IS reducing the metabolic capacity of isolates. It is also possible that the four focal IS-dense sub-lineages may contain additional novel or specialised metabolism that are not accounted for in the current model.

Insertion of IS not only leads to genomic rearrangements as had been observed for SL82 (**Fig. 6**) but have also been shown to impact expression of neighbouring genes via their promoters. One prime example is IS*1* (61, 62). Considering the vast numbers of IS found across these IS-dense sub-lineages and their close proximity to many regulatory, amino acid and carbohydrate metabolism loci (**Fig. S2**), it may be that IS are facilitating metabolic specialisation via provision of a selective advantage for organisms by transcriptionally upregulating remaining metabolic traits in addition to purifying unneeded metabolic traits. For example, a copy of IS*1* was found directly upstream of the *cra* catabolite repressor/activator gene in an SL82 genome (accession GCF_900451215) that regulates the acetolactate *ilvIN* locus responsible for amino acid biosynthesis from pyruvate. Increasing expression *of cra* may provide a selective advantage for SL82 and balance the effects arising from usage loss of 17 core substrates.

The proximity of IS to metabolic genes could be interpreted as: i) IS ferrying these associated metabolic genes with them during insertion; or ii) IS can only insert into metabolically-tolerable sites for the individual cell, where only metabolism that is not essential for survival can be selectively purified. This insertional tolerance model, combined with potential upregulation of these operons is a compelling hypothesis, as this would improve the fitness of an individual strain via metabolic specialisation. This is consistent with the limited examples of directly interrupted, vestigial metabolic loci observed in our study. Future work could entail using transcriptional data to study the impact of insertions within intergenic regions (62).

## Data availability

All data used in this study is available as supplemental material (Figs. S1-4, Tables S1-4). Additionally, all analysis code is available at Figshare (doi: 10.6084/m9.figshare.28341917).

## Author contributions

Conceptualization: KEH, KLW, JH, MMCL

Data Curation: BV, CW

Formal Analysis: BV, CW, HBC, MMCL

Funding Acquisition: MMCL Methodology: BV

Project Administration: KEH, KLW, JH, MMCL

Resources: KEH, KLW, JH, MMCL

Supervision: KEH, KLW, JH, MMCL

Writing – Original Draft Preparation: BV, KLW, MMCL

All co-authors reviewed and approved the submitted manuscript.

## Acknowledgements

This research/work was supported by Monash eResearch capabilities, including M3 and Research Data Storage.

We thank the Institut Pasteur teams for the curation and maintenance of BIGSdb-Pasteur databases at http://bigsdb.pasteur.fr/

## Supplemental material

**Table S1:** Metadata and strain information of all genomes used in this study. Includes BioSample and accession numbers.

**Table S2:** Results of Insertion Sequence analysis on genomes and statistical analyses

**Table S3:** Results of substrate usage predictions using metabolic models and comparisons to phenotypic tests

**Fig. S1:** Scatterplot of number of IS per genome compared to the IS bp:chromosomal bp ratio per genome. Linear regression model fitted to data.

**Fig. S2:** IS families present in each sub-lineage.

**Fig. S3:** Scatterplot comparing IS load and metabolic model features of SL258. Squared polynomial model fitted.

**Fig. S4:** Scatterplot comparing IS load and metabolic model features of all sub-lineages. Squared polynomial model fitted.

