## Supplementary figures and images for "IS-capades of *Klebsiella pneumoniae*: Insertion sequences drive metabolic loss in obscure sub-lineages"

### Figure S1

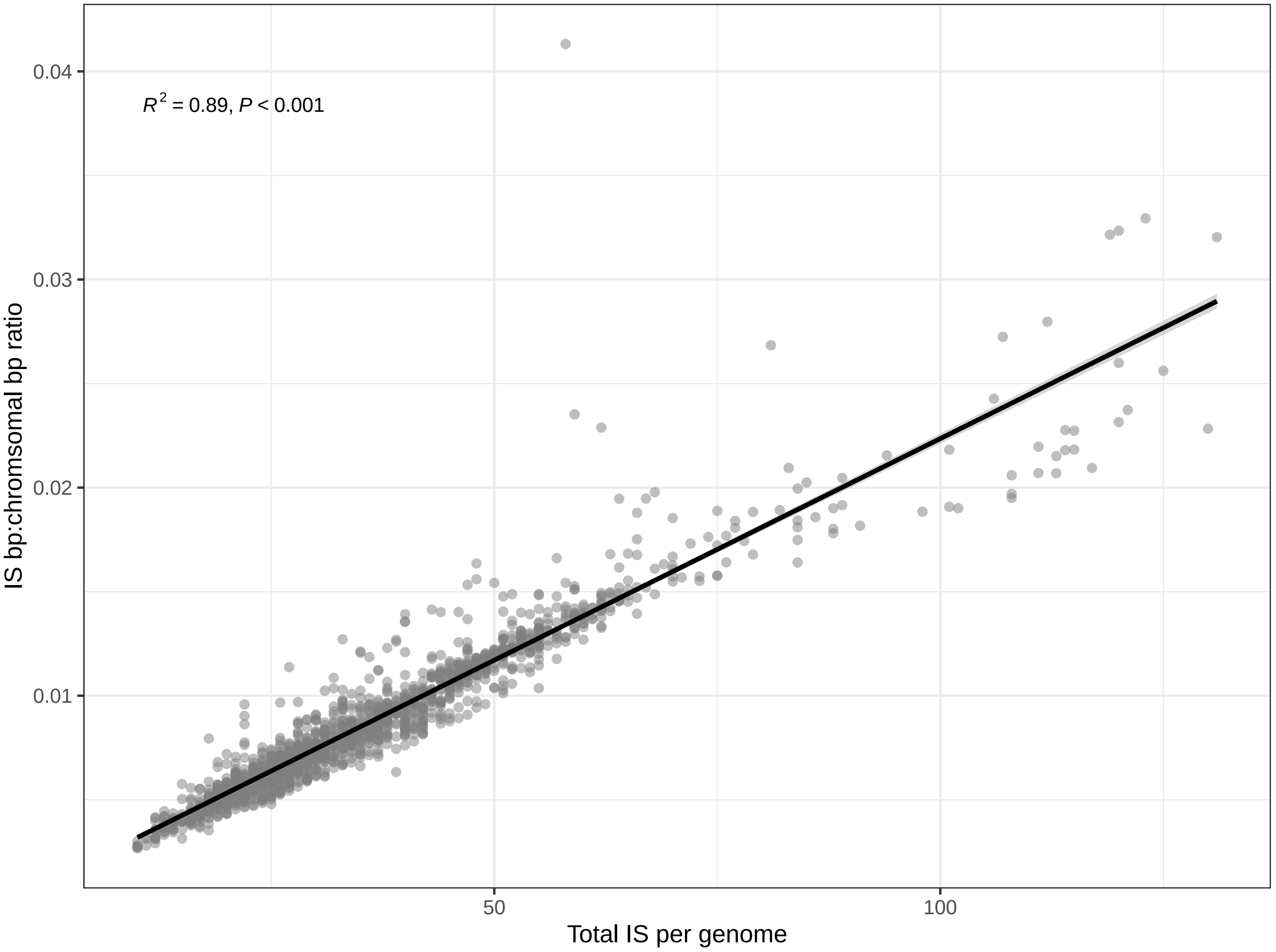

### Figure S2

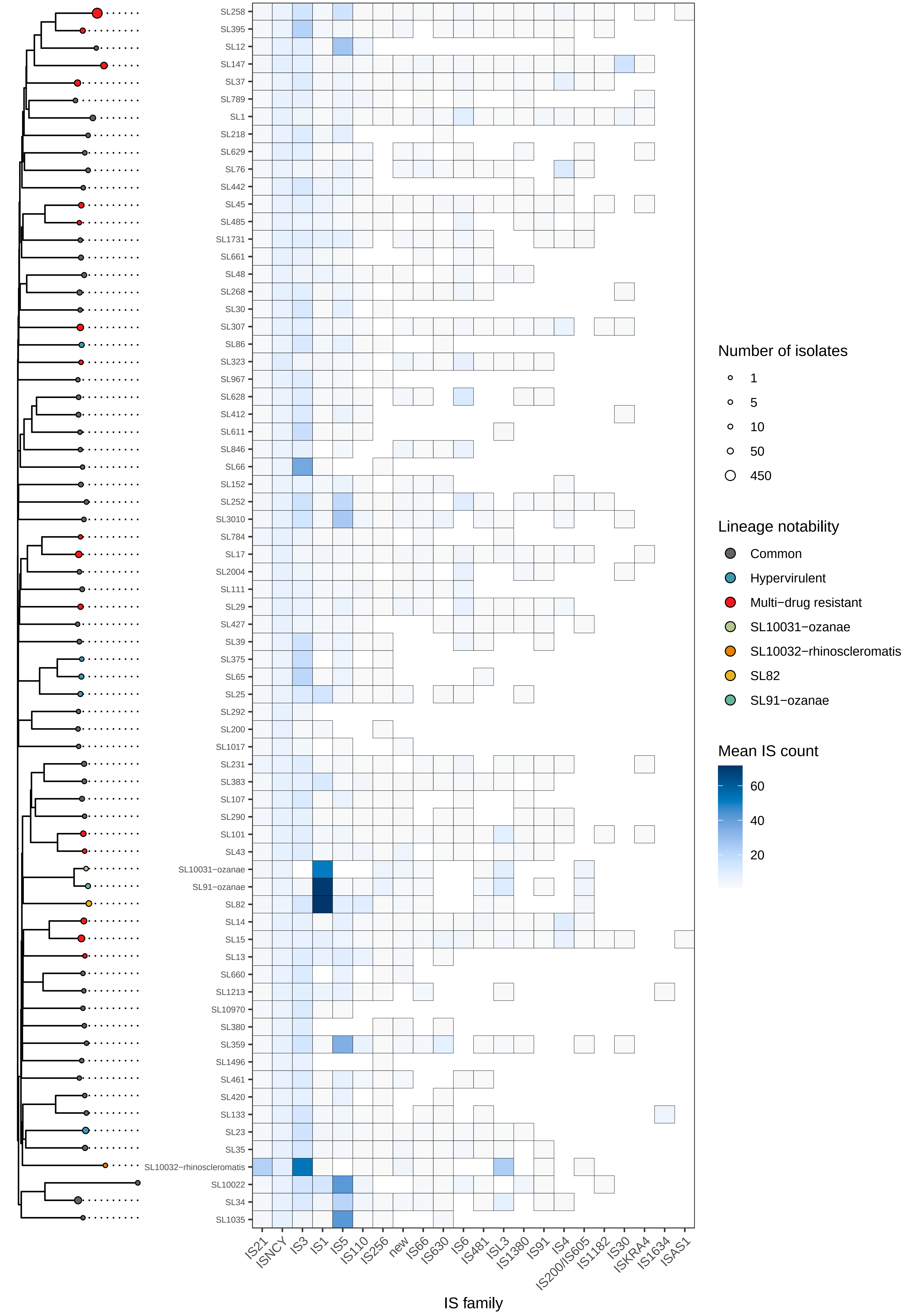

### Figure S3

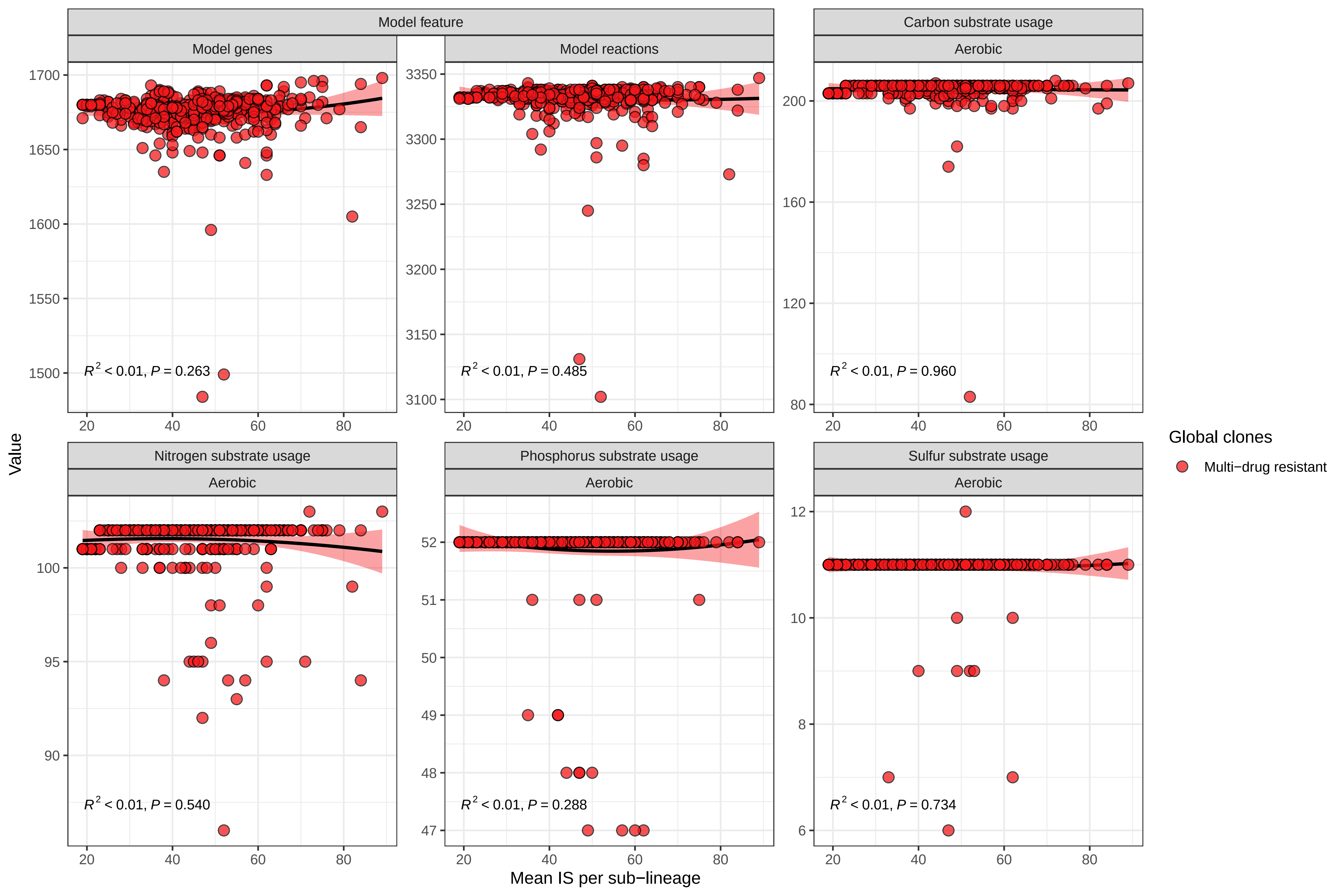

### Figure S4

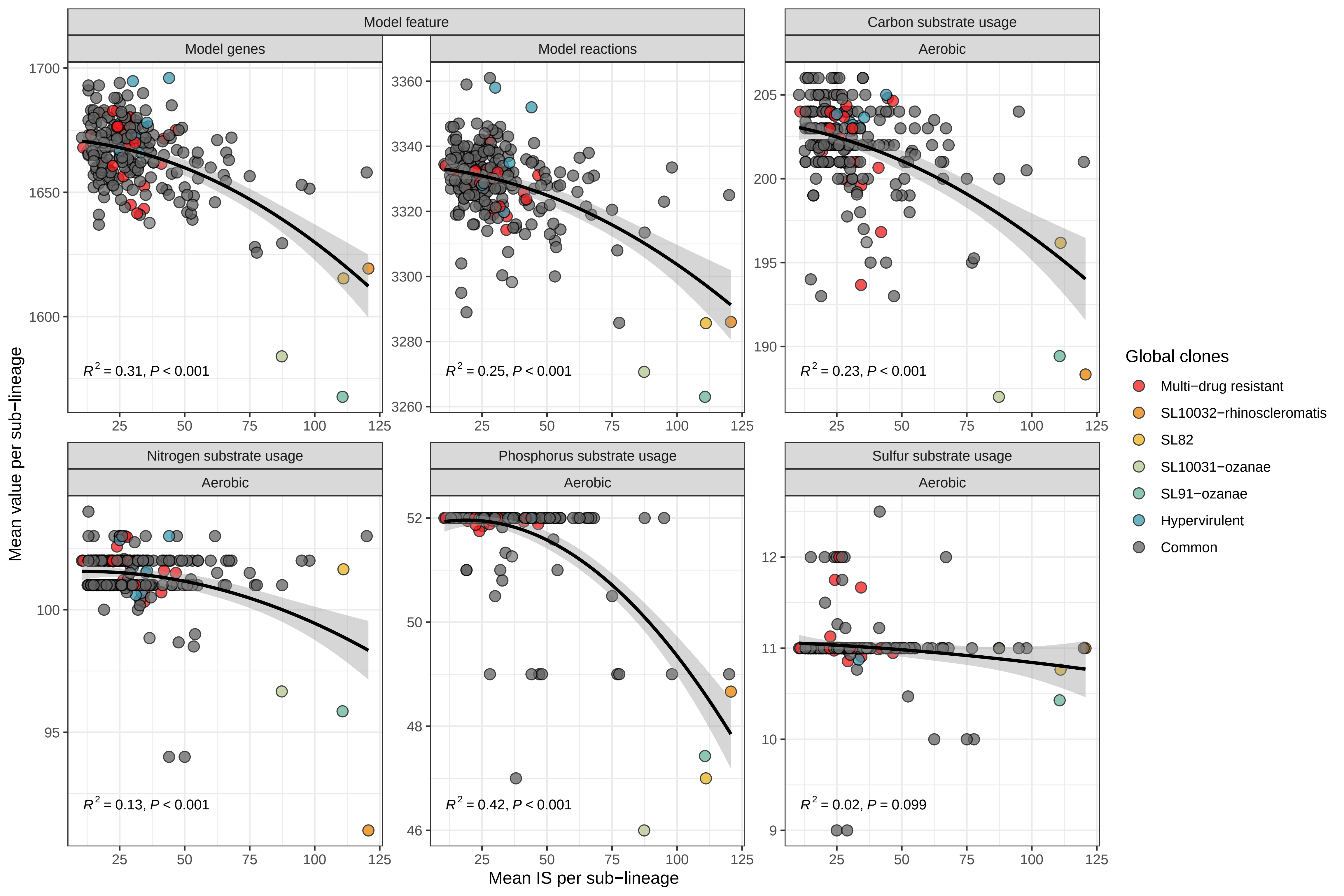
